# Endospores and other lysis-resistant bacteria comprise a widely shared core community within the human microbiota

**DOI:** 10.1101/221713

**Authors:** Sean M. Kearney, Sean M. Gibbons, Mathilde Poyet, Thomas Gurry, Kevin Bullock, Jessica R. Allegretti, Clary B. Clish, Eric J. Alm

## Abstract

Endospore-formers in the human microbiota are well adapted for host-to-host transmission, and an emerging consensus points to their role in determining health and disease states in the gut. The human gut, more than any other environment, encourages the maintenance of endospore formation, with recent culture-based work suggesting that over 50% of genera in the microbiome carry genes attributed to this trait. However, there has been limited work on the ecological role of endospores and other stress-resistant cellular states in the human gut. In fact, there is no data to indicate whether organisms with the genetic potential to form endospores actually form endospores *in situ* and how sporulation varies across individuals and over time. Here, we applied a culture-independent protocol to enrich for endospores and other stress-resistant cells in human feces to identify variation in these states across people and within an individual over time. We see that cells with resistant states are more likely than those without to be shared among multiple individuals, which suggests that these resistant states are particularly adapted for cross-host dissemination. Furthermore, we use untargeted fecal metabolomics in 24 individuals and within a person over time to show that these organisms respond to shared environmental signals, and in particular, dietary fatty acids, that likely mediate colonization of recently disturbed human guts.

## Introduction

To date, there is limited work investigating the relevance of stress-resistant cellular states in the propagation, survival, and function of organisms in the mammalian gastrointestinal (GI) tract. The gut is the only known environment with such a considerable abundance of organisms that form endospores, considered the most stress-resistant of all cell-types (Filippidou et al., 2015). Anaerobic, endospore-forming Firmicutes are numerically dominant members of the GI tract of most animal species (Browne et al., 2016; Ley et al., 2008). Within this group of organisms, the presence of genes for endospore formation suggests that growth in the GI tract favors the maintenance of this large gene repertoire (Browne et al., 2016). The apparent utility of these genes is to allow organisms to enter metabolically dormant states that aid in survival and transmission to new hosts. Passage through the GI tract is likely to trigger sporulation (Angert and Losick, 1998; Flint et al., 2005), but the mechanisms by which this process occurs and the signals that induce sporulation here are mostly unknown, even for well-studied pathogens like *Clostridioides difficile.*

Many endospore-forming organisms in the human gut are in the class Clostridia, the most well-studied of which are the pathogens *C. difficile* and *Clostridium perfringens* (Alexander et al., 1995; Paredes-Sabja et al., 2008). However, Clostridia also includes abundant organisms not known to form endospores, like *Faecalibacterium prausnitzii* (Sokol et al., 2008) and *Roseburia intestinalis* (Png et al., 2010). For *C. difficile,* the role of sporulation is central to disease etiology (Deakin et al., 2012), particularly in patients who experience recurrence. Sporulation and rising levels of antibiotic resistance allow *C. difficile* to persist in the face of antibiotic assault, ensuring that it remains in the environment to rapidly re-colonize its host.

Among Clostridia that do not cause disease, multiple strains of endospore-forming organisms have the capacity to induce T regulatory cells and associated anti-inflammatory cytokines (Atarashi et al., 2011, 2013) involved in sensitivity to, for example, peanut antigen (Stefka et al., 2014). These organisms have recently been shown to provide pathogen resistance in neonatal mice (Kim et al., 2017). Similarly, endospore-forming commensals of the murine GI tract have a central role in mediating the induction of a Th17-type T helper cell response (Ivanov et al., 2009; Kuwahara et al., 2011; Sczesnak et al., 2011). Many Clostridia also produce butyrate as an end-product of metabolism, which can regulate how immune cells interact with gut commensal bacteria (Smith et al., 2013; Furusawa et al., 2013; Van den Abbeele et al., 2013; Eeckhaut et al., 2011; Louis et al., 2010). This group of organisms also boosts production of serotonin by enterochromaffin cells in the intestine, crucial for motility in the gut (Yano et al., 2015). An ecological understanding of sporulation and induction of other resistant states could be informative for how these phenotypes interact with host immunity. For example, such an understanding could inform whether inflammation acts positively or negatively on endospore formation, or whether endospores themselves have immunomodulatory effects.

Resistant cellular states like endospores appear to be adaptive in the mammalian gut environment. It is likely that other non-endospore-forming taxa have evolved analogous resistance strategies for passing between hosts. Persister states may allow non-endospore-forming organisms to enter a metabolically inert state upon exit from the gastrointestinal tract. Toxin-antitoxin systems, associated with persistence in *E. coli,* are overrepresented in Bacteroidetes, Alpha- and Gammaproteobacteria (Makarova et al., 2009)
, and Bacteroidetes are among the most metabolically inactive cells in human fecal samples (Maurice et al., 2013). Further, viable but nonculturable cell (VBNC) states may enable passage through the environment by reducing metabolic needs and affecting cell wall and membrane composition and morphology (Li et al., 2014). These strategies and others not yet studied may play an important role in mediating cross-host bacterial transmission.

Environmental stress resistance protects cells faced with unfavorable conditions. The signals triggering resistance are likely quite varied. Even for well-studied endospore-forming bacteria, inducing sporulation *in vitro* can be difficult, and across strains of one species, signals that induce sporulation in one strain may be insufficient to induce sporulation in others (Kaplan and Williams, 1941). Further, even organisms that abundantly form endospores in their native environment may not do so under conditions permitting vegetative growth. For instance, *Paenibacillus larvae* in honeybees will only form endospores *in vitro* under idiosyncratic conditions designed to mimic the host environment (Dingman and Stahly, 1983). Similarly, certain strains of *Clostridium perfringens* rarely form endospores *in vitro* unless exposed to a specific set of environmental stressors (Kaplan and Williams, 1941). The discrepancy in phenotype of organisms in their native environments compared to *in vitro* argues for culture-free approaches to investigate such phenotypes *in situ.* Enriching for stress-resistant cells in environmental samples provides a means to uncover the actual context in which these states form.

Here, we investigate which organisms are present as endospores or as other resistant cell types in the human gastrointestinal tract. We modified previously described methods to enrich fecal samples for endospores and obtain paired bulk community and resistant fraction 16S rDNA sequence data for 24 healthy individuals and one individual across 24 days. We consistently enriched for putatively endospore-forming taxa in all samples, as well as other taxa, predominantly from the Actinobacteria phylum, that show high levels of lysis resistance. We compared resistant OTUs (rOTUs) and non-resistant OTUs (nOTUs) to identify ecological characteristics differing between these groups. Using a database of rOTUs identified in this study, we find consistent signals for these organisms in their responses to a variety of successional states across multiple independent data sets from prior published studies (Supplementary Table 7). Overall, we show a tight association between the ecological role of these resistant organisms and their distribution within and across human hosts.

## RESULTS AND DISCUSSION

### Sequencing resistant fraction reveals resistant taxa present in human feces

We modified a culture-independent method (Wunderlin et al., 2014) to generate resistant fraction 16S rDNA amplicon data from human feces (Figure 1A). This method entails a series of lysis treatments including heating, lysozyme and proteinase treatment, alkaline and SDS treatment, and hypotonic wash steps followed by DNAse treatment. We extract the resultant resistant fraction pellet alongside the bulk community sample using a bead-beating protocol (see Methods). We validated this method on pure bacterial cultures and endospore cultures prior to treatment of human fecal samples (Supplementary Figure 2) and conducted a small study to validate the reproducibility of the method across samples (Supplementary Table 5). We then proceeded to test the method on feces from a healthy human cohort and from a time series of a single individual.

**Figure 1.**
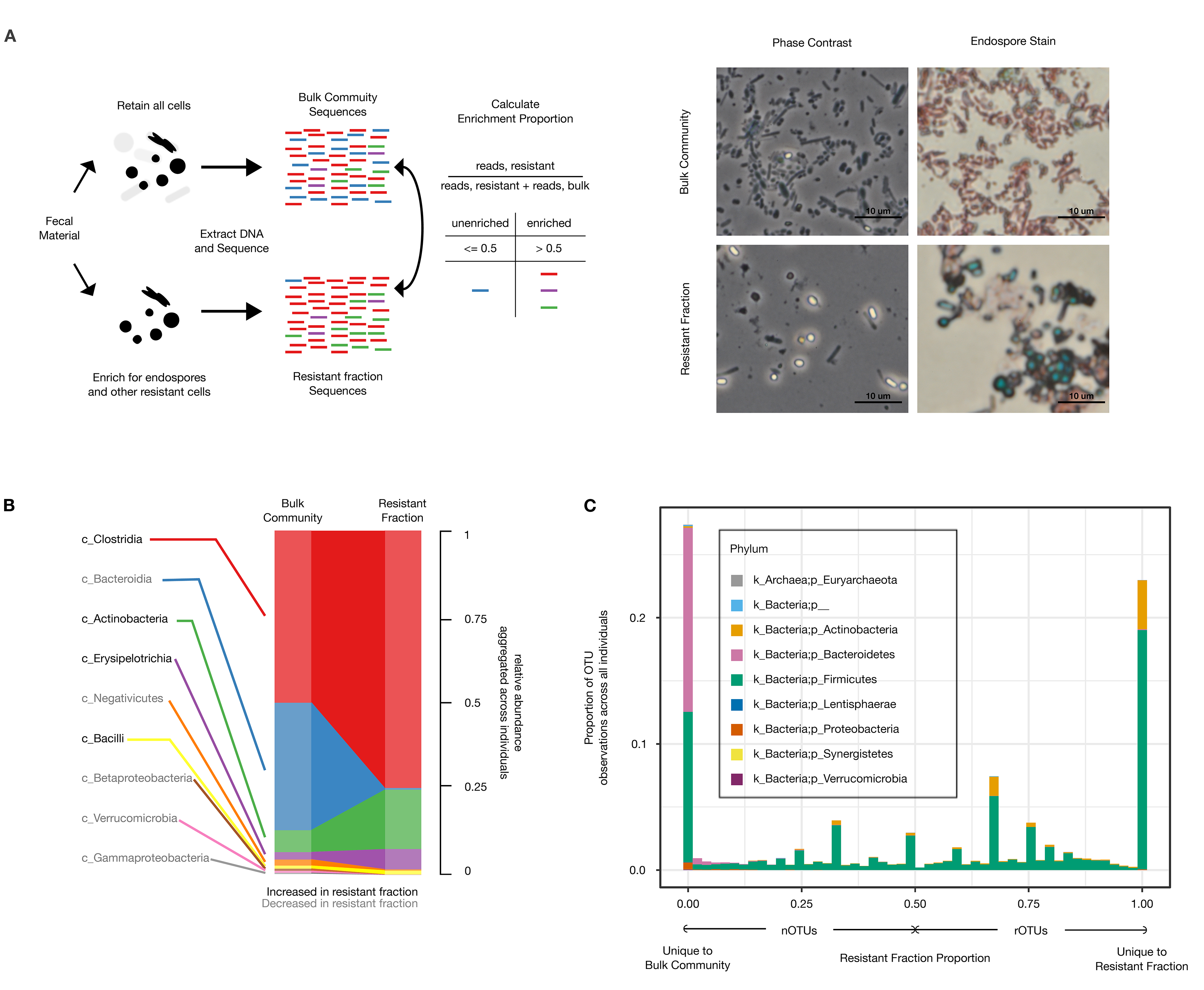
Resistant fraction sequencing of human fecal bacteria. (A) Overview of resistant cell enrichment and 16S rDNA sequencing protocol. Resistant fraction samples are treated with a series of physical, enzymatic, and chemical lysis steps to deplete vegetative cells. DNA from bulk community and resistant fraction samples are extracted via a mechanical lysis protocol, and 16S rDNA libraries prepared. Communities are analyzed to determine the change in abundance of each OTU in the resistant fraction relative to the bulk community. (right) Phase contrast images of bulk community and resistant fraction-phase bright cells are endospores. Endospores stain green when heat fixed with malachite green, vegetative cells appear red from safranin counter stain. (B) Representative results of 16S rDNA profile for bulk community and endospore-enriched samples. Reads from each OTU are summed across 24 individuals to give a meta-bulk and meta-endospore community. Phylogenetic classes in black text increase with resistant fraction; gray text classes decrease with resistant fraction. (C) Distribution of resistant fraction proportion across phyla aggregated across individuals filtered to remove OTUs with single counts in a sample (for visualization purposes). Colors represent phyla. OTUs with a resistant fraction proportion of 0 are absent from the resistant fraction; OTUs with a resistant fraction proportion of 1 are absent from the bulk community and only found in the resistant fraction.

When aggregating the fecal data across our cohort, we see expansion of classes with known endospore-formers in the resistant fraction: Clostridia, Erysipelotrichia, and Bacilli (Figure 1B). We also see depletion of classes lacking endospore-formers (Bacteroidia, Betaproteobacteria, Verrucomicrobia, Gammaproteobacteria). Organisms in the class Actinobacteria were enriched in the resistant fraction, but lack genes considered essential for endospore formation. Although exospore formation is well documented in some families of Actinobacteria (e.g. Actinomycetaceae and Streptomycetaceae), these families have only modest representation in our data. We see high-level resistance primarily from *Bifidobacterium* and *Collinsella,* whose representative genomes lack orthologs for genes thought to be essential to exospore formation.

We suspect that high level resistance in the Actinobacteria is mediated primarily by resistance to lysozyme conferred by cell wall structures common to Actinobacteria (Sekar et al., 2003). Lysozyme is one of the most common and important defense mechanisms used by neutrophils, monocytes, macrophages, and epithelial cells (Fahlgren et al., 2003; Keshav et al., 1991). It is abundant in human milk, a source of *Bifidobacterium* species transferred to breast-feeding infants, and in saliva and mucus, where it serves an antibacterial role (Gueimonde et al., 2007). Attempts to deplete Actinobacteria with achromopeptidase, which has previously been shown to break down Actinobacterial peptidoglycan, had variable efficacy across samples (data not shown). Thus, factors other than cell wall structure may contribute to Actinobacteria resistance.

To quantify the extent of lysis resistance, we calculated the proportion of normalized reads for each sequence variant in the resistant fraction to the sum of its reads in the bulk community and the resistant fraction. We then obtain a finite quantity even for organisms not observed in one of the paired samples. When the proportion exceeds 0.5 we call an OTU enriched in the resistant fraction (Figure 1A). An OTU that is enriched in at least one of the samples in which it is present is considered a resistant OTU (rOTU), and non-resistant (nOTU) otherwise. Using the above definitions, all of the rOTUs are either Firmicutes or Actinobacteria (Figure 1C). In fact, when grouping OTUs at the genus level, the top two most enriched genera *(Bifidobacterium, Collinsella)* are both Actinobacteria.

### Resistant fractions consist of few dominant and many rare OTUs

In order to investigate ecological properties of the resistant cell fraction, we first examined the community structure of resistant fractions and compared these to their bulk community counterparts. After rarefying to the minimum read depth (28639 reads, Supplementary Table 6), we find that resistant fractions are significantly less diverse both in species richness and evenness than their bulk community counterparts (Figure 2A). As we are sampling a subset of the community, this result is not necessarily surprising. However, reduced evenness of the resistant fraction compared to the bulk community suggests dominance of a few organisms coupled with many low abundance organisms. This difference in community structure could entail that resistant fractions are more dissimilar from each other than their bulk community counterparts. Instead, the compositions using Jaccard Distance of resistant fractions tend to be more similar to each other than bulk communities to each other across different people (PERMANOVA p-value < 0.001, Figure 2B). Similarly, using Jensen-Shannon Divergence, resistant fractions cluster together separately from bulk communities implying that resistant fractions may assemble in similar ways across people (Figure 2C). We hypothesized that the coherence between resistant fractions may lead to increased prevalence of organisms found in the resistant fraction across people.

**Figure 2.**
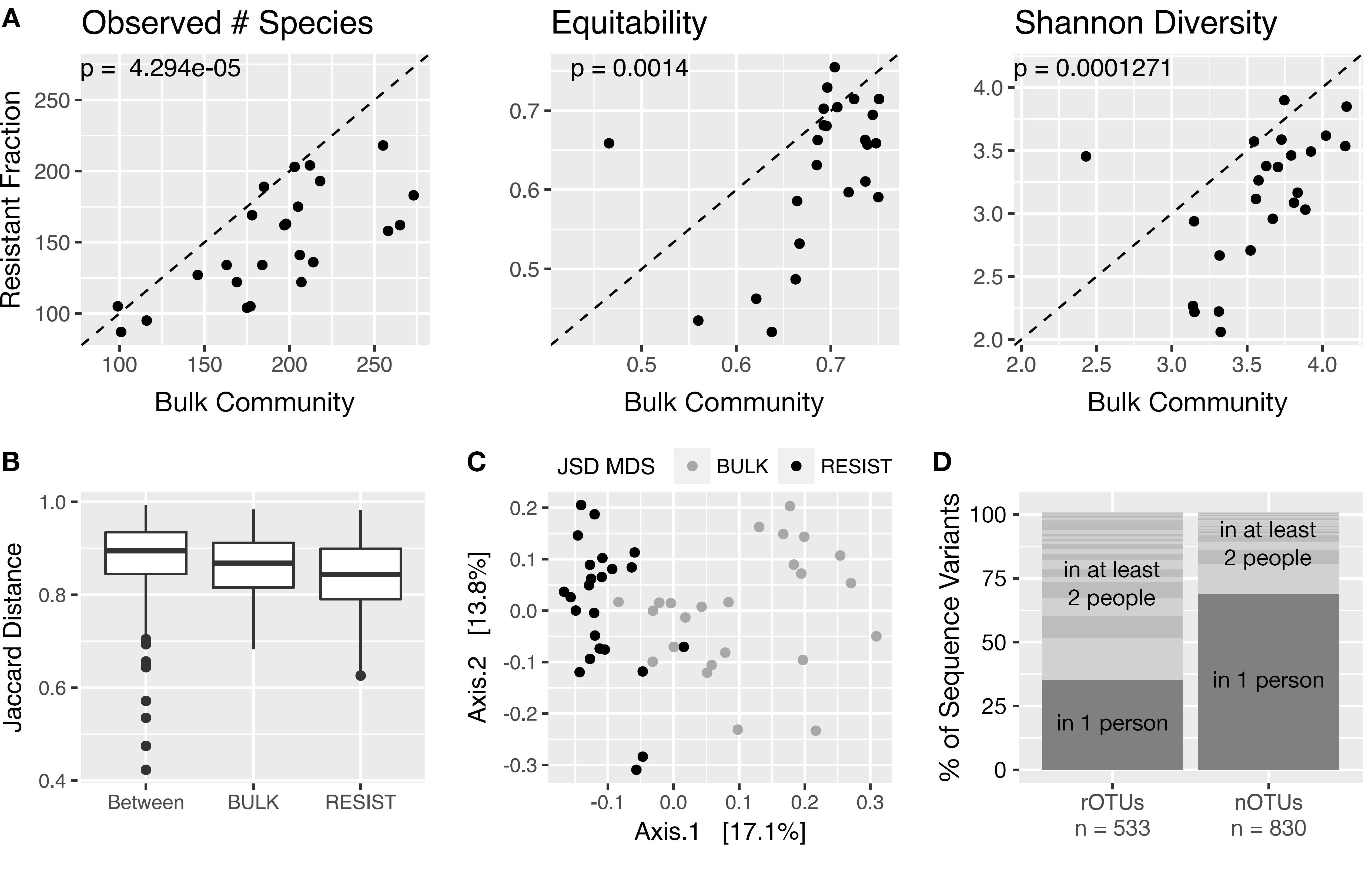
Resistant fraction OTUs are more shared across individuals than bulk community OTUs. (A) Alpha diversity metrics measured for the bulk community (x-axis) and resistant fraction enrichments (y-axis). P-values are for the test of differences between alpha-diversity metric distributions using paired Wilcoxon Rank Sum Test. (B) Distribution of Jaccard distance between resistant fraction and bulk communities, within the resistant fraction, and within the bulk communities. (C) Multidimensional scaling on the Jensen-Shannon Divergence of all resistant fraction and bulk community samples. Black dots represent resistant fraction communities, gray dots represent bulk community samples. (D) Comparison of the number of rOTU sequence variants in only one person and in multiple people to nOTU sequence variants in only one person and in multiple people.

### rOTUs are more shared than nOTUs among individuals

To test the hypothesis that resistant states contribute to prevalence, we examined the frequency with which rOTUs were found among the bulk communities across individuals compared to nOTUs (Figure 2D). First, nOTUs, which are never enriched in the resistant fraction, are significantly less likely to be shared among multiple individuals than rOTUs (Mann Whitney U Test comparing the distribution of the number of individuals sharing each rOTU to the number of individuals sharing each nOTU, p-value = 1e-36). We again see this result by calculating the correlation between the frequency of resistance (the number of times an organism is enriched in the resistant fraction divided by the number of times it is observed) and sharedness (number of individuals an OTU is observed in divided by the total number of individuals), giving a weak, but positive and highly significant correlation (Spearman rho, correlation = 0.23, p-value = 1e-17; Kendall tau, correlation = 0.19, p-value = 1e-17). Finally, when we compare the proportion of rOTUs found in only one person compared to multiple people with that proportion for nOTUs in bulk communities, we find rOTUs are about four times as likely to be found in multiple individuals (Fisher Exact Test, p-value = 1e-33, odds ratio = 4.0).

These results suggests that organisms that do not form resistant states are less likely to be found across multiple individuals than those that do. Yet, rOTUs tend to be less dominant members of the community (median rOTUs = 13.5 counts, median nOTUs = 18 counts, Mann-Whitney U Test, p-value = 0.004). Even though rOTUs are not as dominant as nOTUs, they are more widespread within this cohort.

### Representation of organisms in resistant fractions is heterogeneous across and within individuals

We suspected that the increased prevalence of rOTUs might indicate positive selection on resistance capabilities in this environment. Variation in this trait among related organisms could be indicative of selection. In order to visualize how much of a population is present in a resistant state within a given sample, we scaled 16S rDNA abundance data using V4 16S rDNA qPCR-based estimates of community size (Supplementary Figure 1, Supplementary Table 6, and Supplementary Text) and defined the resistant fraction as the ratio of these scaled reads for each OTU. We plot this quantity on a phylogeny representing 99% OTUs (clustered at 99% nucleotide ID using usearch (Edgar, 2010)) present in at least 8 individuals and up to 24 individuals (Figure 3). First, we note the high variability in the resistant fraction within and across taxa (the average variation is over 50-fold within each taxon). For one Roseburia 99% OTU in particular, this quantity varies over 3 orders of magnitude, suggesting this OTU contains organisms present in a resistant state in some individuals, but not in others.

**Figure 3.**
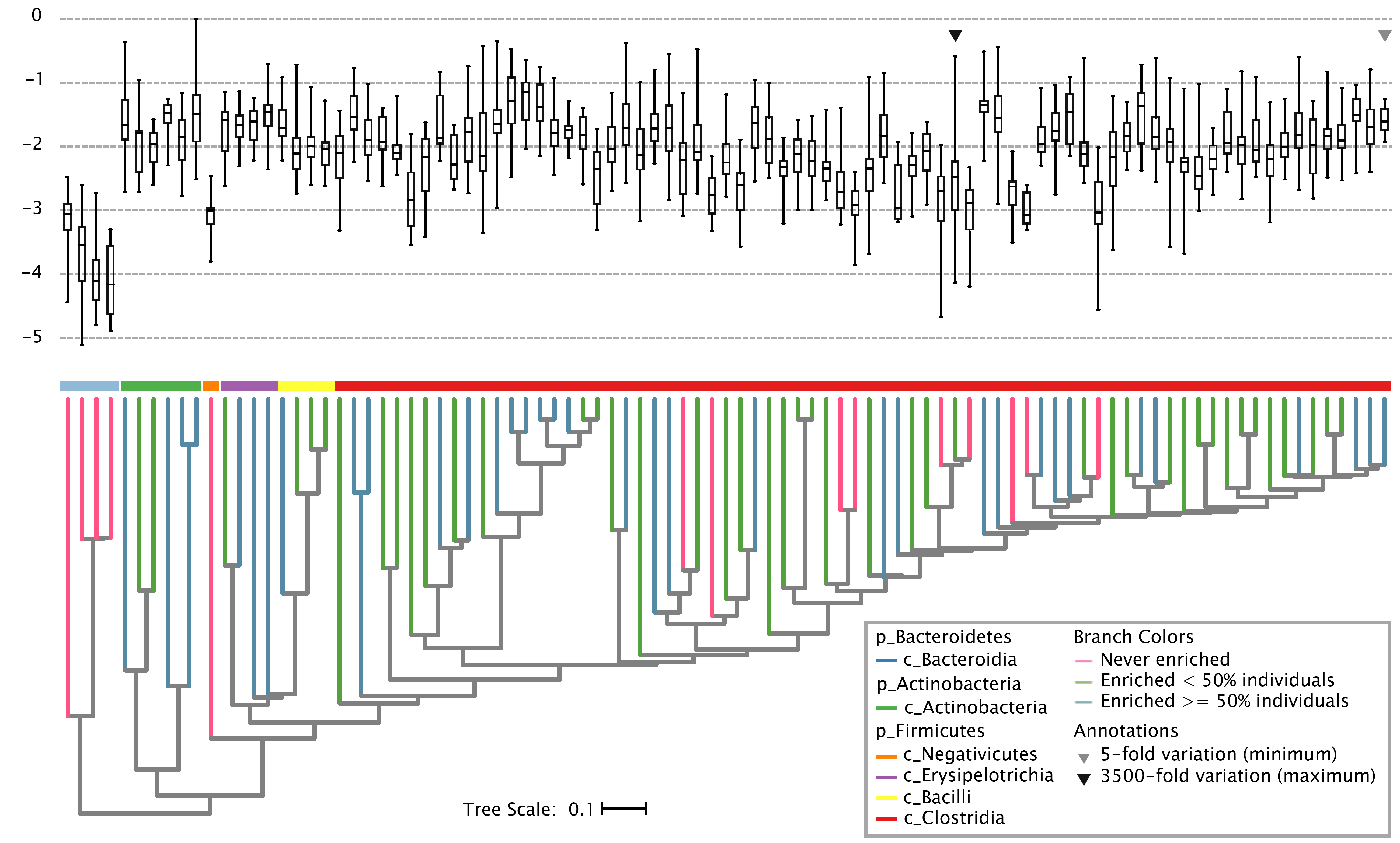
Taxa show heterogeneous patterns of resistant cell fractions across individuals. Phylogenetic placement of the fraction of resistant organisms for taxa present within at least 8 individuals estimated by the ratio of counts scaled by qPCR-estimates of biomass in the resistant fractions and bulk communities. Tree branch colors represent the degree to which a taxonomic group was enriched in the resistant fraction with pink branches never enriched and blue and green branches enriched at least once. Classes within each phylum are shown with a colored bar. Arrows indicate OTUs showing the maximum (black) and minimum (gray) within-OTU variability in enrichment scores.

Furthermore, within a person, OTUs with the same genus classification can be discordant in their degree of resistance. In the individual time series, for example, one *Ruminococcus* 100% OTU is almost always enriched, and another is never enriched (Supplementary Figure 3). The closest matching genomes to these two organisms show differences in sporulation gene content, with the resistant *Ruminococcus* sharing 48/58 core sporulation genes (Galperin et al., 2012), and the non-resistant only 41/58 (Supplementary Figure 4 and Supplementary Tables 1 and 4). We also see that spore maturation proteins *spmA* and *spmB* vary in their presence in genomes of genera with variable enrichment phenotypes. These genes are involved in spore cortex dehydration and heat resistance in *B. subtilis* and *C. perfringens,* so their loss might contribute to differences in the recovery of resistant cells in this work.

The process of entering a resistant state itself might be selected on in this system. There is evidence that the sporulation phenotype is evolving in mammalian guts, as several gut isolates of *B. subitilis* lack genes that negatively regulate sporulation compared to their laboratory counterparts (Serra et al., 2014). Knowing which organisms can form resistant cells in a community does not provide complete information about which organisms do (see supplementary results). Formation of resistant states *in vivo* seems to be highly context dependent. We also note that loss of a single gene (i.e. *spo0A)* in *C. difficile* is sufficient for loss of sporulation, such that retaining endospore formation requires strong purifying selection.

### rOTUs share signals for growth within an individual

Previous evidence has shown that bile acids contribute to outgrowth of *C. difficile* endospores *in vivo* (Francis et al., 2013). As taurocholate is a known germinant for several endospore-forming species (Browne et al., 2016), we wondered whether endospore-formers and resistant organisms more broadly would share dynamic behavior over time, suggesting coherent responses to environmental signals. We compared the Euclidean distance of correlation profiles between organisms to determine whether there were differences between the correlation profiles of rOTUs and nOTUS (Figure 4A). We find that rOTUs are more similar in their correlation profiles than nOTUs (PERMANOVA, p < 0.001). The average correlation between rOTUs in the time series to each other is 0.211 compared to 0.162 for nOTUs to each other (Wilcox rank sum test, p-value = 5e-08): strong correlated behavior in this person associated primarily with rOTUs (Figure 4A). We interpret this result to mean that the dynamic behavior of rOTUs is more strongly coupled: these OTUs respond coherently to environmental signals.

**Figure 4.**
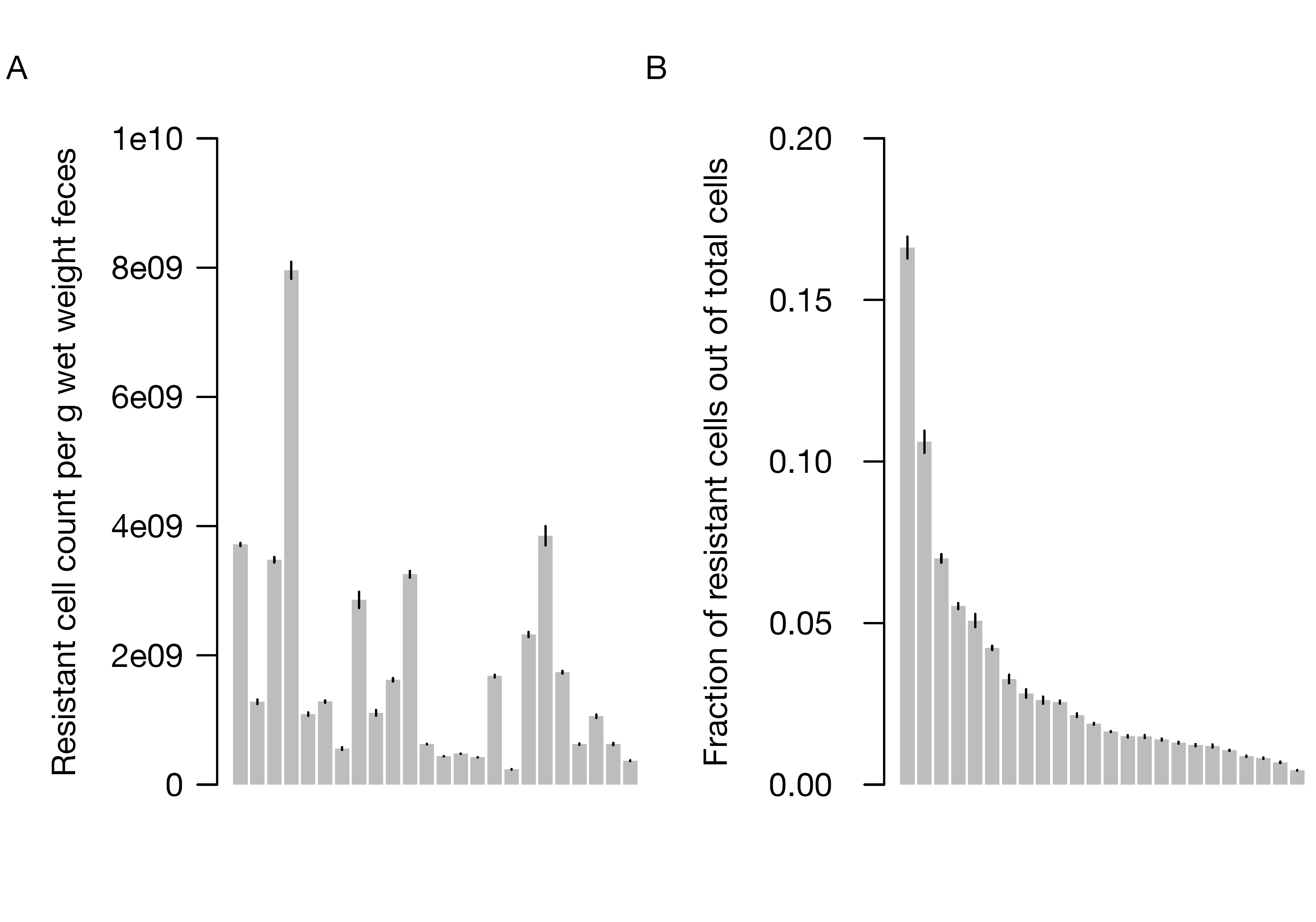
Common signals govern resistant state exit and growth in the GI tract. (A) Boxplot of the distribution of correlation distances (pairwise Euclidean distance between the Spearman correlation vectors for two OTUs) between rOTUs and nOTUs, within rOTUs, and within nOTUs. (B) Abundance of OTUs in the resistant fraction as a function of bile acid exposure for nine phylogenetically distant OTUs.

### rOTUs link growth to fatty acid metabolism

To address whether bile-related signals relate to the dynamics of rOTUs in the time series, we conducted untargeted metabolomics with standards for fatty acid metabolism. We then calculated the Spearman correlations between the median abundance profile of all OTUs clustering with the highly correlated rOTUs and metabolites for which we had standard markers. This cluster tends to correlate positively with long-chain saturated fatty acids, and negatively with long-chain polyunsaturated fatty acids and, notably, taurocholate (Supplementary Table 2). An OTU in the genus *Bilophila,* known to use taurocholate for sulfite reduction (Devkota et al., 2012) also clusters with these organisms, and shows a strong relationship to markers of milkfat consumption (see supplementary text). We suspect that taurocholate metabolism by members of this group drives down the concentration of taurocholate in stool. Additionally, saturated fatty acid concentration in the stool measures fatty acids escaping absorption in the small intestine. This process would be negatively impacted by microbial metabolism of taurocholate, as it more efficiently emulsifies saturated fats than glycine-conjugated primary bile acids (Devkota et al., 2012). Fecal concentrations of taurocholate reflect secretion of unmetabolized taurocholate, which should increase if taurocholate metabolism by the gut microbiota decreases.

### Resistant cells lose resistance in response to physiological bile acid concentrations

As a more direct test of the coupling of rOTUs to bile acid concentration, we dosed ethanol-treated feces (to kill vegetative cells without the additional harshness of the resistant fraction DNA enrichment protocol) with increasing concentrations of bovine bile in aqueous solution. We then measured the depletion of OTUs from the endospore-enrichment using 16S rDNA sequencing (Figure 4B). When correcting for biomass via qPCR, nearly 20% of OTUs identified in the resistant fraction apparently germinated in response to bile acids (log-link quasipoisson generalized linear model, p-value < 0.05, Supplementary Table 3). The true fraction of resistant cells that lose resistance in response to bile acids is likely higher, as many endospores require an activation step (i.e. heating at 80°C or treatment with lysozyme as for *C. difficile* (Sorg and Sonenshein, 2010)) before they will respond to germinants.

Notably, most ethanol-resistant OTUs began to show a germination-like response at 0.5% bile (Figure 4B), which is near the concentrations found in the human small intestine (Ceuppens et al., 2012). Although Clostridia and other putative endospore-formers make up the majority of organisms that lose resistance in response to bile acids, genera in the Actinobacteria and other resistant cells also show this response when approaching physiological concentrations. These conserved responses suggest that the same cues can mediate loss of resistance in distantly related organisms, similar to the conserved resuscitation response of dormant bacteria to peptidoglycan(Shah et al., 2008).

### rOTUs exhibit shared dynamics in diverse contexts

Correlated behavior, increased prevalence, and shared signals for growth among rOTUs indicated that these organisms might exhibit a global response during disturbances of various kinds. To test this hypothesis, we made a sequence database of rOTUs within our cohort, and used this database to identify putative rOTUs in other datasets (Figure 5A). While certainly not all rOTUs in our dataset will map to organisms forming resistant states in other datasets, we assume that some strains or species within an rOTU are capable of forming a resistant state at some time.

**Figure 5.**
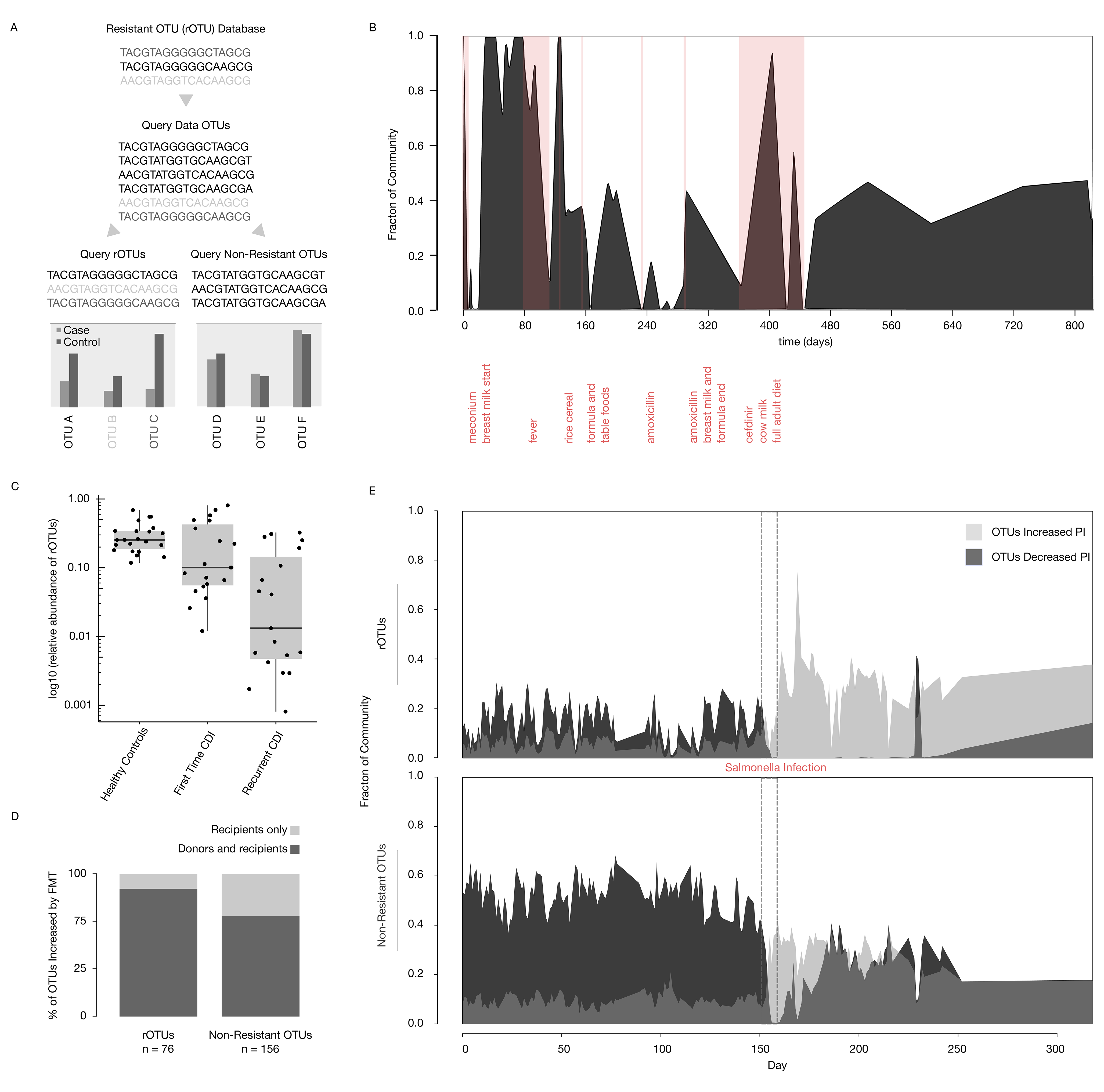
Resistant OTUs show disproportionate turnover in diverse contexts. (A)Overview of approach for identifying resistant-cell forming OTUs in 16S rDNA sequencing datasets. rOTU database sequences are matched to sequences in other datasets, and then patterns within those datasets among the identified rOTUs are determined. (B) Fraction of rOTUs present during microbial colonization of an infant gut annotated with major diet and health perturbations. rOTUs encompass both putative endospore-forming organisms and those not known to form endospores, but which possess a resistant state (Actinobacteria and non-endospore-forming Firmicutes) (C) Fraction of rOTUs present as a function of *C. difficile* infection status (fCDI = first time *C. difficile* diagnosis, rCDI = at least 3 episodes of *C. difficile* infection following initial treatment) (D) Fraction of rOTUs and all other OTUs (non-resistant OTUs) transferred from donors to recipients by fecal microbiota transplant. (E) Time series of rOTUs (top) and all other (non-resistant) OTUs (bottom) from a human male infected with Salmonella, with OTUs significantly more abundant pre-infection (dark gray) and significantly more abundant post-infection (light gray).

We expected that increased prevalence and shared signals for growth would lead to enhanced colonization of the developing infant gut microbiota (Koenig et al., 2011). The lysozyme-resistant members of the Actinobacteria and Bacillales dominate the infant gut microbiota for most of the first 80 days of life and do not equilibrate until the infant starts a full adult diet (Figure 5B). Early colonization by these rOTUs connects a resistant state to development of the infant gut microbiome. Here, lysozyme resistance might be essential for semi-selective transmission of *Bifidobacterium,* as human breast milk is rich in lysozyme (Chandan et al., 1964), potentially lysing non-resistant cells. Others have shown endospore-formers negatively associate with vertical transmission from mother to infant (Nayfach et al., 2016), but other environmentally resistant states as in the Actinobacteria may be important for vertical transmission as well.

Depletion of endospore-forming clades is common during infection with *C. difficile.* We predicted a strong signal for rOTUs in individuals infected with *C. difficile,* due to its sporulation requirement for transmission (Deakin et al., 2012). We find a significant depletion of rOTUs dependent on *C. difficile* infection status (Figure 5C), with a serial depletion of rOTUs from healthy to first time diagnosis to recurrent patients (Allegretti et al., 2016). Because of this depletion in rOTUs, we expected that fecal microbiota transplant (FMT) might transfer relatively more rOTUs than other OTUs (Youngster et al., 2014). Indeed, among OTUs shared with donors, 90% of rOTUs increase in abundance following FMT, compared to 77% for the rest of the community (Fisher exact test, p-value = 0.008) (Figure 5D).

We suspected that rOTUs are a particularly malleable component of the microbiota. To test this hypothesis, we measured the turnover of rOTUs in the time series of an otherwise healthy male individual who was infected by Salmonella (David et al., 2014).

New rOTUs almost completely replaced old rOTUs following this perturbation. By contrast, fewer OTUs from the rest of the community were lost and gained. This result holds both when examining the number of OTUs replaced (Fisher exact test, p-value = 6e-12) as well as the change in abundance of these OTUs (Figure 5E). We see again that rOTUs exhibit coherent responses to changes in the gut environment, most pronounced in systems with dramatic perturbations. Colonization of newly vacant niches favors rOTUs, likely transmitted in an endospore or other resistant state to germinate in an environment replete with nutrients (including untransformed bile acids). In the absence of a fully functioning microbiota, rOTUs appear to fill open niches more readily than nOTUs.

## CONCLUSION

Gut bacteria in the resistant fraction were more shared across individuals and showed more correlated dynamics compared to non-resistant organisms. Resistant taxa show greater turnover following large-scale disturbance events, as in the case of *C. difficile* and *Salmonella* infection, which suggests that many of these organisms are sensitive to environmental fluctuations and respond to stress by entering into a dormant, seed-like state. Environmental sensitivity and high turnover rates of resistant taxa provide an opportunity to manipulate the composition of the human gut microbiota through targeted perturbations and replacements. Because of the therapeutic relevance of Clostridia endospores (Atarashi et al., 2011, 2013; Kim et al., 2017; Stefka et al., 2014; Yano et al., 2015), determining the exact conditions that permit their replacement may be of high value for future microbiota-based therapeutics. Here, we found that the growth of many resistant organisms was associated with dietary fatty acids. If this result extends to more individuals, one can imagine a therapeutic strategy coupling dietary changes with introduced resistant cells to enable robust colonization and engraftment.

## MATERIALS AND METHODS

### Contact for reagent and resource sharing

Further information may be obtained from the Lead Contact Eric J. Alm (; address: Massachussetts Institute of Technology Cambridge, MA, 02139, USA)

### Experimental Model and Subject Details

#### Human Subjects

Human subject enrollment and sample collection was approved by the Institutional Review Board of the Massachusetts Institute of Technology (IRB Approval Number: 1510271631). Informed consent was obtained from all subjects. 12 male and 12 female healthy human subjects (age range 21-65) with no history of antibiotic use in the last six months were enrolled in the study. In total, 24 fecal samples were collected from these individuals and an additional 24 fecal samples were collected from one male individual (age 24) over 24 days for culturing and DNA isolation.

### Method Details

#### Fecal Sample Processing and Storage

Fecal samples were collected and processed in a biosafety cabinet within 30 minutes of defecation. Samples (5 g) were suspended in 20 mL of 1% sodium hexametaphosphate solution (a flocculant) in order to bring biomass into solution as described previously (Wunderlin et al., 2014). Fecal samples were bump vortexed with glass beads to homogenize, and centrifuged at 50 × g for 5 min at room temperature to sediment particulate matter and beads. Triplicate aliquots of 1 mL of the supernatant liquid were transferred into cryovials and stored at −80° C until processing. For the time series, samples were collected at approximately 24-hour intervals.

#### Resistant Fraction Enrichment from Fecal Samples

We modified a previously published method (Wunderlin et al., 2014) for endospore sequencing to increase throughput and decrease signal from contaminating, non-endospore forming organisms. Fecal samples previously frozen at −80° C were thawed at 4° C prior to use, and 500 μL was aliquoted for resistant fraction, while the remaining 500 μL was saved for bulk community DNA extraction. Samples were centrifuged at 4° C and 10,000 × g for 5 minutes, washed and then resuspended in 1 mL Tris-EDTA pH 7.6. Samples were heated at 65° C for 30 minutes with shaking at 100 rpm and then cooled on ice for 5 minutes. Lysozyme (10 mg/mL) was added to a final concentration of 2
mg/mL and the samples were incubated at 37° C for 30 minutes with shaking at 100 rpm. At 30 minutes, 50 uL Proteinase K (>600 mAU/ml) (Qiagen) was added and the samples incubated for an additional 30 minutes at 37° C. Next, 200 uL 6% SDS, 0.3 N NaOH solution was added and the samples incubated for 1 hour at room temperature with shaking at 100 rpm. Samples were then centrifuged at 10,000 rpm for 30 minutes. At this step, a pellet containing resistant endospores should be visible or slightly visible in the sample, and the pellet is washed three times at 10,000 × g with 1 mL chilled sterile ddH2O. The pellet is then resuspended in 100 uL ddH2O, and treated with 2 uL DNAse I (Ambion) to remove residual contaminating DNA with incubation at 37° C for 30 min. The DNAse is killed by addition of 10 μL Proteinase K (Qiagen) and incubation at 50° C for 15 minutes, followed by incubation at 70° C for 10 minutes to inactivate Proteinase K. At this step, microscopic examination of samples is used to confirm the presence of phase-bright (or phase-dark) spores. The sample is then ready for downstream extraction and sequencing.

#### Extraction of Nucleic Acids

We extracted DNA from both the original sample suspended in sodium hexametaphosphate and the output of the resistant fraction. Both the original sample and the resistant fraction were extracted with MoBio PowerSoil Isolation Kit (MoBio Laboratories, Inc.) with three 10 minute bead-beating steps followed by sequential collection of ⅓ of the solution to enhance recovery of endospore DNA as shown previously (Bueche et al., 2013). DNA was extracted from bacterial pure cultures, fecal enrichment cultures, and endospores using the same protocol as for fecal samples. DNA from bacterial colonies for 16S rDNA Sanger sequencing confirmation or qPCR was obtained by homogenizing colonies in alkaline polyethylene glycol buffer as described previously (Chomczynski and Rymaszewski, 2006).

#### 16S rDNA Library Preparation and Sequencing

Libraries for paired-end Illumina sequencing were constructed using a two-step 16S rRNA PCR amplicon approach as described previously with minor modifications (Preheim et al., 2013). In order to account for cross-sample and buffer contamination, triplicate negative controls comprising resistant fraction extraction blanks, nucleic acid extraction blanks, and PCR negatives were included during library preparation and samples were randomized across the plate. The first-step primers (PE16S_V4_U515_F, 5′ ACACG ACGCT CTTCC GATCT YRYRG TGCCA GCMGC CGCGG TAA-3′; PE16S_V4_E786_R, 5′-CGGCA TTCCT GCTGA ACCGC TCTTC CGATC TGGAC TACHV GGGTW TCTAA T 3′) contain primers U515F and E786R targeting the V4 region of the 16S rRNA gene, as described previously (Preheim et al., 2013). Additionally, a complexity region in the forward primer (5-YRYR-3′) was added to help the image-processing software used to detect distinct clusters during Illumina next-generation sequencing. A second-step priming site is also present in both the forward (5′-ACACG ACGCT CTTCC GATCT-3′) and reverse (5′-CGGCA TTCCT GCTGA ACCGC TCTTC CGATC T-3′) first-step primers. The second-step primers incorporate the Illumina adapter sequences and a 9-bp barcode for library recognition (PE-III-PCR-F, 5′-AATGA TACGG CGACC ACCGA GATCT ACACT CTTTC CCTAC ACGAC GCTCT TCCGA TCT 3′; PE-III-PCR-001-096, 5′-CAAGC AGAAG ACGGC ATACG AGATN NNNNN NNNCG GTCTC GGCAT TCCTG CTGAA CCGCT CTTCC GATCT 3′, where N indicates the presence of a unique barcode.

Real-time qPCR before the first-step PCR was done to ensure uniform amplification and avoid overcycling all templates. Both real-time and first-step PCRs were done similarly to the manufacturer's protocol for Phusion polymerase (New England BioLabs, Ipswich, MA). For qPCR, reactions were assembled into 20 μL reaction volumes containing the following: DNA-free H2O, 8.9 μL, HF buffer, 4 μL, dNTPs 0.4 μL, PE16S_V4_U515_F (3 μM), 2 μL, PE16S_V4_E786_R (3 μM) 2 μL, BSA (20 mg/mL), 0.5 μL, EvaGreen (20X),1 μL, Phusion, 0.2 μL, and template DNA, 1 μL. Reactions were cycled for 40 cycles with the following conditions: 98° C for 2 min (initial denaturation), 40 cycles of 98 C for 30 s (denaturation), 52° C for 30 s (annealing), and 72° C for 30s (extension). Samples were diluted based on qPCR amplification to the level of the most dilute sample, and amplified to the maximum number of cycles needed for PCR amplification of the most dilute sample (18 cycles, maximally, with no more than 8 cycles of second step PCR). For first step PCR, reactions were scaled (EvaGreen dye excluded, water increased) and divided into three 25-μl replicate reactions during both first- and second-step cycling reactions and cleaned after the first-and second-step using Agencourt AMPure XP-PCR purification (Beckman Coulter, Brea, CA) according to manufacturer instructions. Second-step PCR contained the following: DNA-free H_2_O, 10.65 μL, HF buffer, 5 μL, dNTPs 0.5 μL, PE-III-PCR-F (3 μM), 3.3 μL, PE-III-PCR-XXX (3 μM) 3.3 μL, Phusion, 0.25 μL, and first-step PCR DNA, 2 μL. Reactions were cycled for 10 cycles with the following conditions: 98° C for 30 s (initial denaturation), 10 cycles of 98° C for 30 s (denaturation), 83° C for 30 s (annealing), and 72° C for 30s (extension). Following second-step clean-up, product quality was verified by DNA gel electrophoresis and sample DNA concentrations determined using Quant-iT PicoGreen dsDNA Assay Kit (Thermo Fisher Scientific). The libraries were multiplexed together and sequenced using the paired-end with 250-bp paired end reads approach on the MiSeq Illumina sequencing machine at the BioMicro Center (Massachusetts Institute of Technology, Cambridge, MA).

#### qPCR

For testing of the resistant fraction protocol, qPCR was carried out as described in the **16S rDNA Library Preparation and Sequencing** section. Total bacterial abundance was quantified using the same primers. For quantification of Firmicutes and Actinobacteria, primer sequences were obtained from (Fierer et al., 2005). Primers were used at the same concentrations as 16S primers, and annealing temperatures were adjusted to the appropriate temperature for the corresponding primer pairs.

#### 16S rDNA Sequence Data Processing and Quality Control

Paired-end reads were joined with PEAR (Zhang et al., 2014) using default settings. After read joining, the complexity region between the adapters and the primer along with the primer sequence and adapters were removed. Except where specified otherwise, sequences were processed using the DADA2 (Callahan et al., 2016) pipeline in R, trimming sequences to 240 bp long after quality filtering (quality trim Q10) with maximum expected errors set to 1. A sequence variant table was generated using DADA2. Sequence variants were classified using RDP (Maidak et al., 1996; Wang et al., 2007). The resulting count tables were used as input for analysis within R.

#### Identifying High Confidence Endospore-Forming & Resistant OTUs

We developed a workflow for identifying organisms showing increased abundance in the resistant fraction relative to the bulk community. First sequences present at more than 1% in negative control samples were removed from the DADA2 sequence variant table. The resultant pruned sequence variant table was down-sampled to the minimum read depth (25808) and then used to calculate a resistance score for each sequence variant in each sample as Resistance Score = (# of reads in resistant fraction)/(# of reads in resistant fraction + # of reads in bulk community). We then identified sequence variants that had an resistance score greater than 0.5 (more reads in the resistant fraction than in the bulk) at least once across samples, denoting these sequence variants rOTUs. All other OTUs were considered nOTUs. Next, because there were several sequence variants found in the resistant fraction that were absent from all bulk communities (291 with 0 prevalence of 795 total rOTUs), we excluded calculations (as in Figure 4D) involving these OTUs, which would have apparently deflated prevalence estimate from the bulk community samples.

To compile a list of high-confidence resistant fraction-enriched organisms, we took a similar strategy as before, but also included OTUs which had 0 counts in the bulk community but non-zero counts in the resistant fraction. The OTUs increased in abundance in the resistant fraction compared to the bulk community in more than half of the samples present (excluding singletons) were included in this list.

#### Genomic Spore Gene Content

Protein sequences in *Bacillus subtilis subtilis 168* from genes identified as shared among all spore-forming Bacilli and Clostridia(Galperin et al., 2012) were downloaded from UniProt (http://www.uniprot.org/) to make a spore gene database. All genomes as of August 2016 from 9 genera of the Clostridia in containing OTUs that were both significantly enriched at times in the resistant fraction and significantly unenriched were downloaded from NCBI. A standard tblastn approach was used to identify homologues in the downloaded genomes with the corresponding genes in the spore gene database. After identifying presence/absence of spore genes, genome spore gene profiles were hierarchically clustered using UPGMA on the binary distance (Jaccard) matrix.

#### Metabolite profiling

Metabolites were measured using liquid chromatography tandem mass spectrometry (LC-MS) method operated on a Nexera X2 U-HPLC (Shimadzu Scientific Instruments; Marlborough, MA) coupled to a Q Exactive hybrid quadrupole orbitrap mass spectrometer (Thermo Fisher Scientific; Waltham, MA) methods. Stool samples (200mg/mL in 1% sodium hexametaphosphate) were homogenized using a TissueLyser II(Qiagen). Stool homogenates (30 μL) were extracted using 90 μL of methanol containing PGE2-d4 as an internal standard (Cayman Chemical Co.; Ann Arbor, MI) and centrifuged (10 min, 10,000 x g, 4°C). The supernatants (2 μL) were injected onto a 150 x 2.1 mm ACQUITY UPLC BEH C18 column (Waters; Milford, MA). The column was eluted isocratically at a flow rate: 450μL/min with 20% mobile phase A (0.1% formic acid in water) for 3 minutes followed by a linear gradient to 100% mobile phase B (acetonitrile with 0.1% formic acid) over 12 minutes. MS analyses were carried out using electrospray ionization in the negative ion mode using full scan analysis over *m/z* 70-850 at 70,000 resolution and 3 Hz data acquisition rate. Additional MS settings were: ion spray voltage, −3.5 kV; capillary temperature, 320°C; probe heater temperature, 300 °C; sheath gas, 45; auxiliary gas, 10; and S-lens RF level 60. Raw data were processed using TraceFinder 3.3 (Thermo Fisher Scientific; Waltham, MA) and Progenesis QI (Nonlinear Dynamics; Newcastle upon Tyne, UK) software for detection and integration of LC-MS peaks.

#### Bile germination tests

Treatment of fecal samples with ethanol has previously been shown to allow culture-based recovery of endospore-forming organisms (Browne et al., 2016). To this end, fresh fecal samples were homogenized in 50% ethanol (250 mg/mL), incubated for 1 hour under aerobic conditions with shaking at 100 rpm, and washed three times (5 min, 10,000 x g) with sterile water to remove residual ethanol. Serial dilutions from 1e-4-10% (w/v) bile bovine oxgall (Sigma) were prepared in sterile water and 2.5 mL ethanol-treated fecal suspension mixed in triplicate with 2.5 mL each of these bile solutions. Samples were incubated under aerobic conditions for 2 hours at 37° C with 200 rpm shaking, and then transferred to −80° C prior to resistant fraction extraction and 16S rDNA library preparation.

#### Bile germination analysis

We transformed 16S rDNA sequencing counts generated by the bile germination tests again using the cumulative sum-scaling transformation (Paulson et al., 2013). Under the assumption that cells in the resistant fraction can only decrease or remain the same during treatment, we searched for negative relationships between bile acid concentration and abundance that would indicate and OTU had germinated. To identify significant negative relationships, we first fit a generalized linear model (GLM) with a log-link quasi-Poisson distribution to the normalized counts of OTUs present in the control sample with bile acid concentration as the predictor variable. We then identified the OTU with the strongest positive trend in the data (that with the highest positive slope and lowest p-value). We assume that OTUs increase due only to compositional effects (that is, this OTU has not germinated but its abundance apparently increases due to loss of other OTUs), and we use the slope estimated from the fit of this model to detrend the other dose-response data so as to constrain the abundance of this apparently increasing OTU to be constant. We do so by dividing counts of all OTUs by exp(slope*bile acid concentration), which is also a measure of the depletion of the endospore-enrichment biomass. From this detrended dose-response data, we again fit a quasipoisson GLM and identify putatively germinating OTUs as those having a significant (p < 0.05) negative slope.

#### Analysis of Infant Gut Time Series

SRA files containing 16S rDNA Sequences were downloaded from Genbank under accession no. SRA012472) (Koenig et al., 2011). Sequences were generated using a Roche 454 pyrosequencer. In order to simplify analysis of the dataset, these sequences were again processed using the protocol outlined for processing of the original dataset in this paper. However, sequences were quality trimmed using Q20 to 230 base pairs, and the retained sequences were used to call 100% OTUs. OTUs were assigned taxonomies using RDP and 100% OTUs were collapsed into taxonomic names. As very few sequences matched between datasets when using uclust, these taxonomic names were instead used to identify organisms as potential resistant cell-formers based on the correspondence to the RDP-assigned taxonomic names of high confidence resistant cell-formers identified previously. While this approach loses information given the noted heterogeneity in resistance phenotypes even among closely related strains, the original sequences themselves are still proxies for having this phenotype, and so the results of such analysis must be interpreted keeping this observation in mind.

The relative abundance of organisms identified in the infant gut time series as putative resistant-cell formers were summed, and the dynamics of this resistant cell-forming population in the infant gut was visualized over time.

#### Analysis of 16S rDNA sequence files from first time and recurrent *C. difficile* infection

The open reference 97% OTU table including RDP taxonomic annotations from Allegretti et al 2016 was used for this analysis (Allegretti et al., 2016). OTU IDs were mapped using uclust to the corresponding genus level OTUs identified as rOTUs from this study. Patients were grouped either as healthy, first-time *C. difficile* infection (fCDI), or recurrent *C. diffiicle infection* (rCDI), and the fraction of rOTUs was calculated by summing their relative abundances within each patient. A Mann Whitney U test was used to determine whether there were significant differences in the total relative abundance of rOTUs across groups with a Bonferroni multiple hypothesis test correction.

#### Analysis of 16S rDNA sequence files from fecal microbiota transplant in relapsing *C. difficile* infection

This dataset was obtained from (Youngster et al., 2014). To simplify analysis, an existing closed-reference GreenGenes 97% OTU table generated by the original authors was used. Closed-reference OTU IDs were mapped back to GreenGenes reference sequences, and sequences were assigned to the resistant cell-former database sequences again using uclust as for the adult time series.

Unique pre-FMT, post-FMT, and donor samples were separated in the dataset. We again identified organisms that had significantly different relative abundance (Benjamini-Hochberg adjusted Mann-Whitney U test p < 0.05) across the groups for our analysis. We again obtained four categories of OTUs: nonresistant and resistant cell-formers that were elevated in the donor and the post-FMT samples relative to the pre-FMT samples. We used the Fisher exact test on the contingency table containing the number of OTUs in each of the previously mentioned categories to identify whether OTU engraftment from the donor was different across the groupings.

#### Analysis of 16S rDNA sequence files in adult time series pre-and post-Salmonella Infection

Illumina HiSeq sequencing files containing 16S rDNA sequences from the stool of a healthy adult male (David et al., 2014) were downloaded and processed as described for the original dataset in this paper, except that sequences were trimmed to 101 base pairs as described previously before calling 100% OTUs due to the use of shorter read sequencing technology. Sequences were assigned to the resistant cell-former database sequences using uclust constrained with the parameters: --id 99 –usersort –libonly, in order that sequences from this dataset would be assigned only to resistant cell-formers.

In order to assess the presence of differential turnover between resistant and non-resistant cell formers in this dataset, we identified organisms that had significantly different relative abundance (Benjamini-Hochberg adjusted Mann-Whitney U test p < 0.05) before Salmonella infection starting at day 151 (days 0-150) and after the end of infection at day 159 (days 160-252). We partitioned these OTUs into four sets for our analysis: non-resistant and resistant cell formers whose median abundance was higher post-infection and those whose median abundance was lower post-infection. We used the Fisher exact test on the contingency table containing the number of OTUs in each of the previously mentioned categories to identify whether the OTU turnover was different across the groupings.

## DATA AVAILABILITY STATEMENT

All amplicon sequencing data generated in this study have been can be accessed on the US National Center for Biotechnology Information SRA database under BioProject PRJNA389431. Metabolomic data and DADA2 sequence variant tables are available online through Github (https://github.com/microbetrainer/Spores).

## CODE AVAILABILITY STATEMENT

All custom scripts generated in R to analyze the data in this paper will be made available through GitHub (https://github.com/microbetrainer/Spores).

## ACKNOWLEDGMENTS

We thank Fatima Hussain and Mathieu Groussin for extensive discussion and experimental advice. We thank the MIT BioMicro Center for sequencing service. Funding was provided by the Broad Institute BN10 Training Grants. SM Kearney was funded by an NSF Graduate Research Fellowship.

## SUPPLEMENTARY INFORMATION

Supplementary information is available at ISME’s website

---

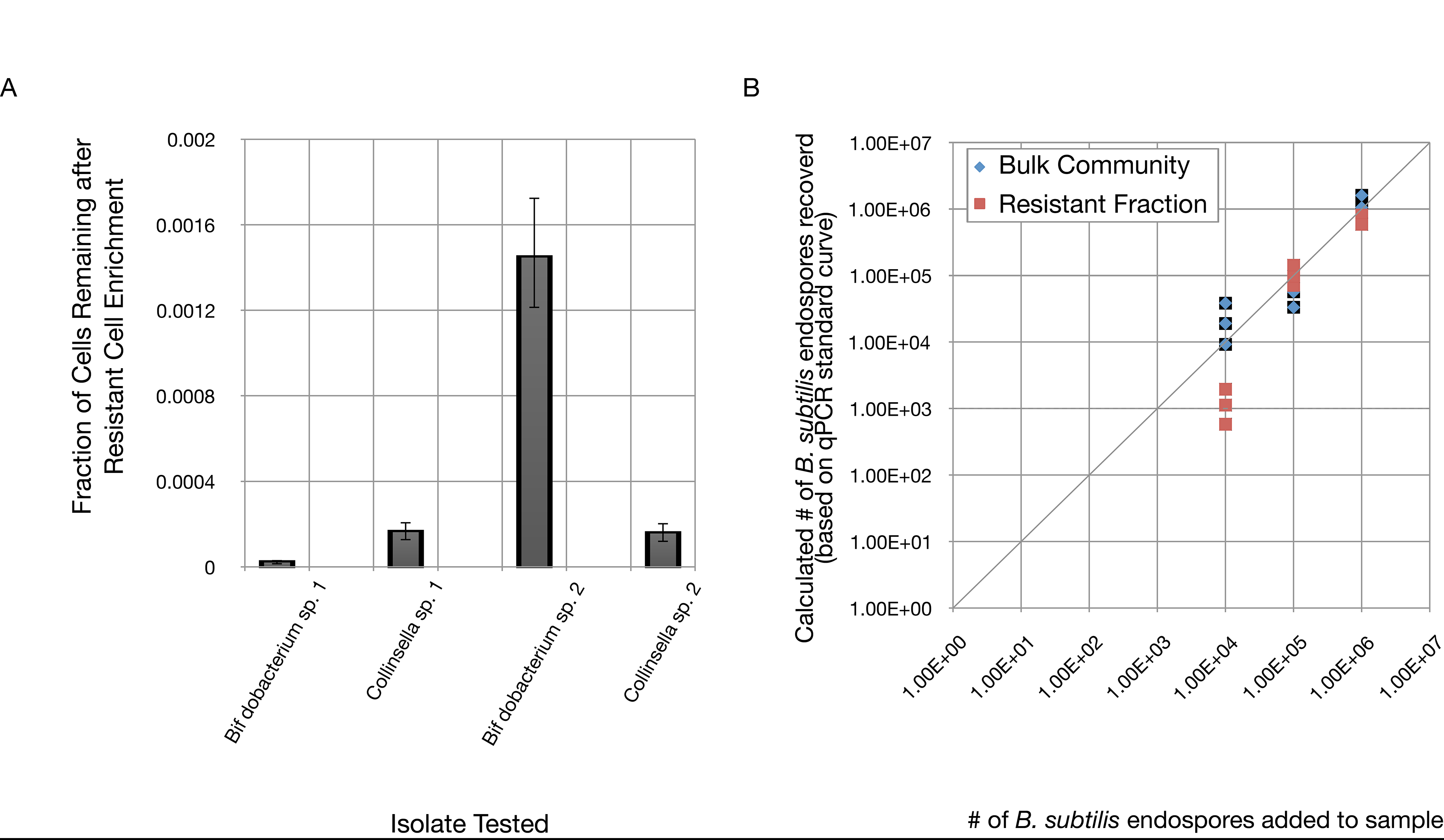

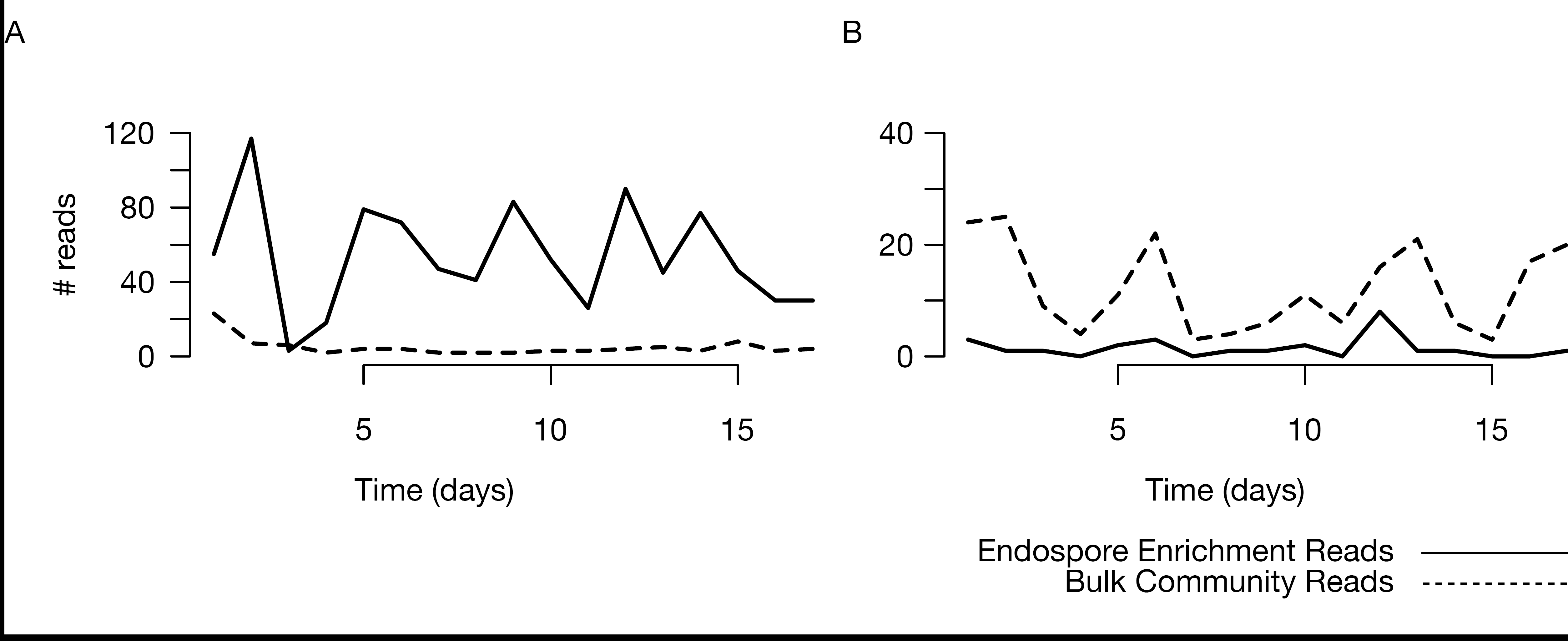

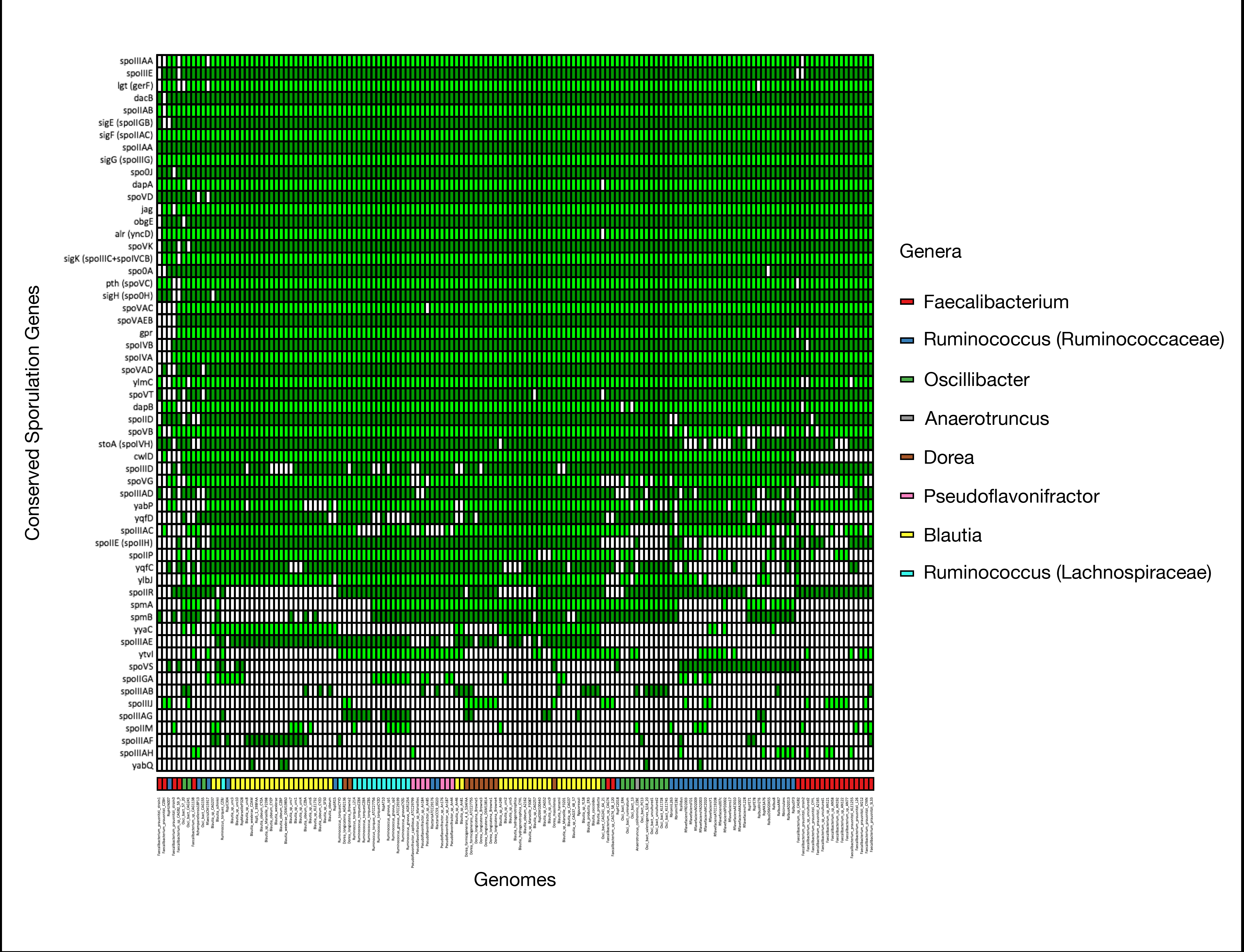

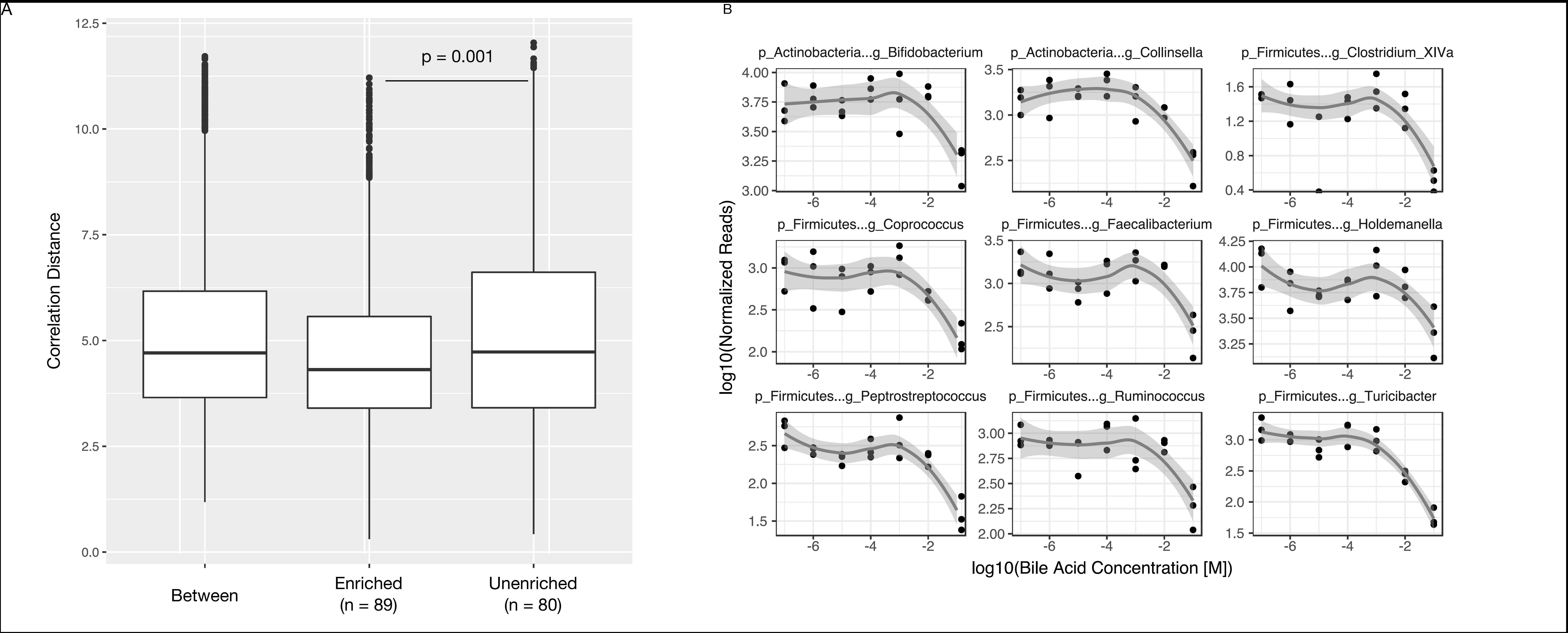

